# Functional characterization of Arabidopsis ARGONAUTE 3 in reproductive tissue

**DOI:** 10.1101/500769

**Authors:** Pauline E. Jullien, Stefan Grob, Antonin Marchais, Nathan Pumplin, Clement Chevalier, Caroline Otto, Gregory Schott, Olivier Voinnet

**Affiliations:** Institute of Molecular Plant Biology, Swiss Federal Institute of Technology Zurich (ETH-Zurich), Zurich, Switzerland; Institute of Plant Sciences, University of Bern, Bern, Switzerland; Department of Plant and Microbial Biology, University of Zurich and Zurich-Basel Plant Science Center, University of Zurich, Zurich, Switzerland

**Keywords:** Seeds, *Arabidopsis*, small RNA, Argonautes, AGO3, reproduction

## Abstract

*Arabidopsis* encodes ten ARGONAUTE (AGO) effectors of RNA silencing, canonically loaded with either 21-22nt small RNAs (sRNA) to mediate post-transcriptional-gene-silencing (PTGS) or 24nt sRNAs to promote RNA-directed-DNA-methylation. Using full-locus constructs, we characterized the expression, biochemical properties, and possible modes of action of AGO3. Although *AGO3* arose from a recent duplication at the *AGO2* locus, their expression differs drastically, with AGO2 being expressed in both male and female gametes whereas AGO3 accumulates in aerial vascular terminations and specifically in chalazal seed integuments. Accordingly, *AGO3* down-regulation alters gene expression in siliques. Similar to AGO2, AGO3 binds sRNAs with a strong 5’-adenosine bias, but unlike most *Arabidopsis* AGOs - AGO2 included - it binds efficiently both 24nt and 21nt sRNAs. AGO3 immunoprecipitation experiments in siliques revealed that these sRNAs mostly correspond to genes and intergenic regions. in a manner reflecting their respective accumulation from their loci-of-origin. AGO3 localizes to the cytoplasm and co-fractionates with polysomes to possibly mediate PTGS via translation inhibition.

**Significance statement:** The regulation of gene expression by small RNAs is key for proper plant development and defense. Here, we characterize Arabidopsis AGO3 expression pattern, microRNA regulation and biochemical properties during sexual reproduction.

## INTRODUCTION

RNA silencing is an ancient mechanism found in plants, animals, and fungi that controls endogenous gene expression and fends off invasive nucleic acids including viruses and transposable elements. RNA silencing relies on the production of small RNAs (sRNAs) (Bologna and Voinnet, 2014) by DICER-LIKE RNase-III enzymes (DCL) cleaving double-stranded RNA (dsRNA) precursors. sRNAs are loaded into effector proteins called Argonautes (AGOs), which mediate sequence-specific post-transcriptional gene silencing (PTGS) at the RNA level, or transcriptional gene silencing (TGS) at the chromatin level. The loading specificity for AGO proteins relies, at least partly, on the length and identity of the 5’ terminal nucleotide of the sRNAs (Mi *et al.*, 2008).

The *Arabidopsis* genome encodes ten *AGOs*, which are phylogenetically divided into three distinct clades, namely *AGO4-6-8-9, AG01-5-10*, and *AGO2-3-7*, indicating potential functional redundancy within these clades (Mallory and Vaucheret, 2010). The *AGO4-6-8-9* clade is responsible for RNA-directed-DNA-Methylation (RdDM) and its members are collectively referred to as the “RdDM AGOs”. AGO4, the most ubiquitous and best-studied member of this clade, recruits the DNA methyltransferase DRM2 to target loci to catalyse cytosine methylation (Zilberman *et al.*, 2003; Law and Jacobsen, 2010; Zhong *et al.*, 2014). AGO6 acts in partial redundancy with AGO4 but is less ubiquitously expressed and is able to target RdDM via loading of 24nt sRNAs or, under certain circumstances, 21-22nt sRNAs (Zheng *et al.*, 2007; Havecker *et al.*, 2010; Mccue *et al.*, 2014). AGO9 has a specific role during reproduction (Olmedo-Monfil *et al.*, 2010) whereas *AGO8* is thought to be a pseudogene (Takeda *et al.*, 2008). The *AGO1-5-10* clade is involved in PTGS and is mainly linked to micro RNAs (miRNAs) (Zhu *et al.*, 2011; Bologna and Voinnet, 2014). AGO1, the main member of this clade loads predominantly miRNAs in healthy plants and is ubiquitously expressed (Bologna and Voinnet, 2014). As a consequence, *ago1* mutants display strong pleiotropic phenotypes. AGO10 regulates shoot apical meristem development by binding miR165/166 (Zhu *et al.*, 2011; Liu *et al.*, 2009). The AGO5 wild-type function remains unclear, but an *ago5* dominant mutant allele prevents female gametophyte development (Tucker *et al.*, 2012).

The last clade, comprising *AGO2-3-7* seems to have more specialized functions. AGO7 is involved in the trans-acting (ta)siRNA pathway: loaded with miR390, it targets non-coding TAS3 transcripts for cleavage, resulting in production of secondary siRNAs that regulate the expression of AUXIN RESPONSE FACTORs (ARF3 and ARF4), which are important for the establishment of leaf polarity (Fahlgren *et al.*, 2006; Montgomery *et al.*, 2008). Like AGO1, Arabidopsis AGO2 serves a role in antiviral silencing against *Turnip crinkle virus* (TCV) and many other viruses (Harvey *et al.*, 2011; Jaubert *et al.*, 2011; Carbonell *et al.*, 2012; Pumplin and Voinnet, 2013). AGO2 is also involved in resistance against the phytopathogenic bacterium *Pseudomonas syringae* by binding the miR393 passenger strand (miR393*) to regulate PR1 protein secretion (Zhang *et al.*, 2011). AGO2 has been shown to regulate Plantacyanin by binding miR403 (Maunoury and Vaucheret, 2011). Finally, AGO2 has been also implicated in double-stand break repair upon genotoxic treatments (Wei *et al.*, 2012). Interestingly, in contrast to the PTGS involvement of other member of its clade, AGO3 has been described to preferentially bind 5’Adenosine 24 nt transposon-related sRNAs upon salt stress (Zhang *et al.*, 2016). AGO3 mis-expression could partially complement an *ago4* mutation, suggesting a role in the TGS pathway rather than PTGS.

Despite some data being available under salt-stress, the expression pattern, sRNA loading capacity, potential roles and mode(s) of action of AGO3 in native conditions have eluded investigation so far. In this study, we conducted a functional characterization of *Arabidopsis* AGO3, which, together with its closest homolog AGO2, represent a unique clade among the plant AGOs containing the PIWI-domain catalytic triad DDD motif. *AGO3* arose from a recent duplication event at the *AGO2* locus, resulting in highly similar proteins but unrelated promoter sequences that cause distinct and specific expression patterns, including, in reproductive tissues, the main focus of our study.

## RESULTS

### *AGO2* and *AGO3* arose from a recent duplication

*AGO2* and *AGO3* are directly in tandem on *Arabidopsis* chromosome 1. To investigate the locus structure in detail, we generated a dot-plot representation of the locus self-alignment (Figure 1a). This led to the identification of a duplication break-point in the first exon within an *AtCOPIA* transposon-containing region (*AtCopia27-ATlTE36140* in *AGO2* and *AtCopia28-ATlTE36160* in *AGO3*;(Buisine *et al.*, 2008)). The corresponding region is particularly rich in glycine and arginine (34%G and 12%R in *AGO3* first exon) and thus, referred to as “glycine-rich-repeat” or GRR. The GRR itself is duplicated in *AGO3* compared to *AGO2*, giving rise to a significantly longer exon 1 (949bp in *AGO3* as opposed to 421bp in *AGO2*). The remaining exon 2 and 3, coding for the conserved PAZ and PIWI domains, display 74.16% amino acid identity between AGO3 and AGO2 opposed to only 38.12% with AGO1 and 29.6% with AGO4.

**Figure 1.**
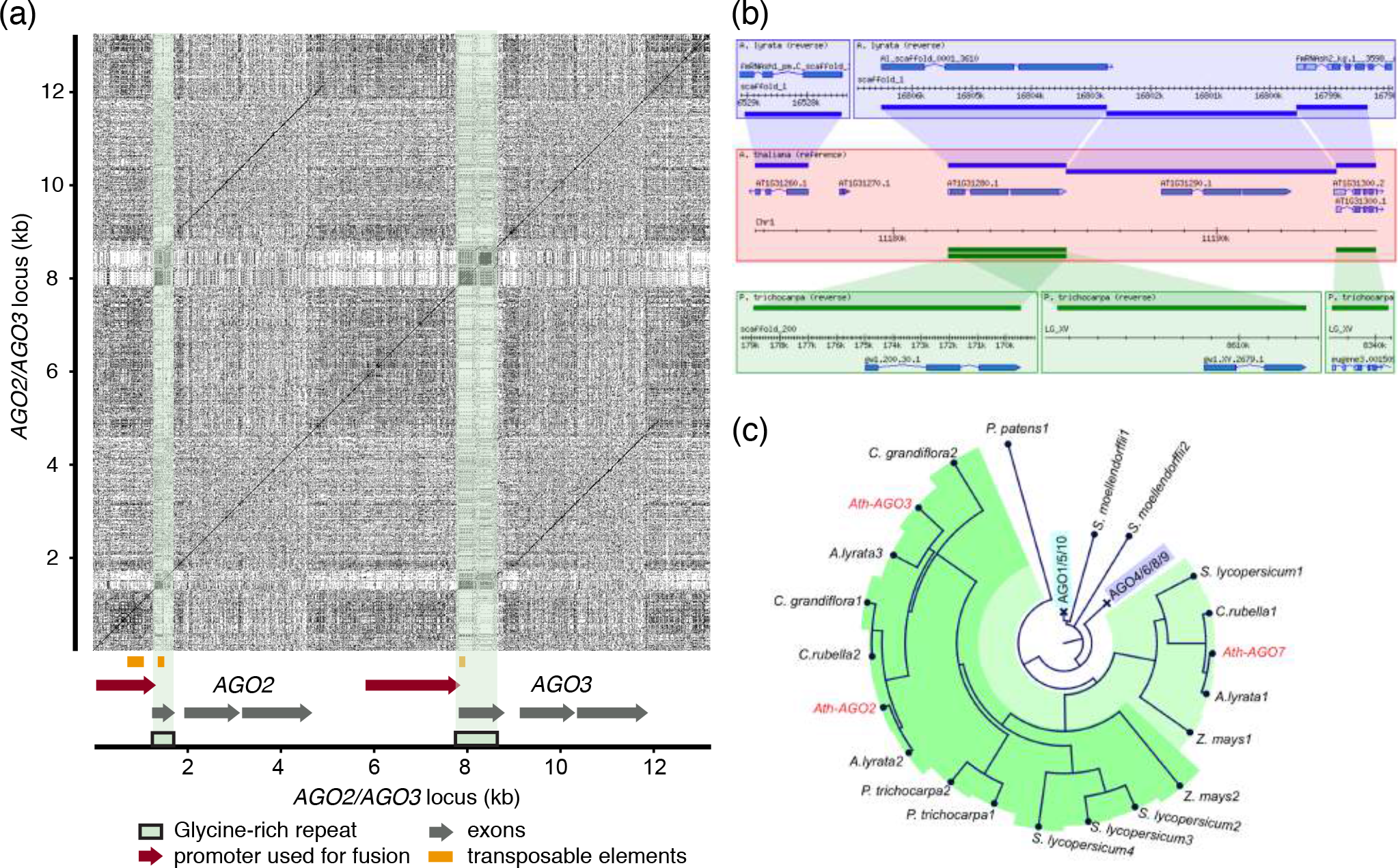
*AGO3* and *AGO2* arose from a recent duplication of the coding region. (a) Dot plot visualization of the alignment of the *AGO2/AGO3* locus to itself showing that the duplication happens at the Glycine rich repeats and comprises the coding sequence but not the promoter. (b) Snap shot of Generic Synteny Browser (TAIR synteny viewer) showing synteny of the *A.thaliana* (*Ath*) *AGO2/AGO3* locus with *A.lyrata* but not with *P.trichocarpa*. (c) Circular phylogram showing that the duplication exists in the *Arabidopsis* lineage but not in in *Capsella* lineage (*C.rubella* and *C.grandiflora*). The AGO1/5/10 and AGO4/6/8/9 branches are collapsed.

To trace the origin of the *AGO2/AGO3* duplication, we first analyzed the syntenic organization of the locus (McKay *et al.*, 2010), which is conserved between *A. thaliana* and *A.lyrata*, but not with *P. trichocarpa*, suggesting a recent duplication event (Figure 1b). Further phylogenic analyses using putative AGOs from various plant species (Figures 1c and S1) revealed that the duplication occurred within the *Arabidopsis* lineage, since the *AGO2/AGO3* pair is present in *Arabidopsis* spp. (*A. thaliana* and *A. lyrata*) but absent in the closely-related *Capsella rubella*.

The amino acids of the PIWI-domain catalytic triad (DDD) required for RNA endonucleolytic cleavage are conserved between AGO2 and AGO3 but differ from the DDH triad found in all other *Arabidopsis* AGOs (Figure S2a). The presence of this catalytic triad and the previous findings that AGO2 is slicing-proficient, suggest that AGO3 also has endonucleolytic cleavage capacities (Poulsen *et al.*, 2013; Carbonell *et al.*, 2012). Close inspection of the sequence alignments of AGOs from the AGO2/AGO3 clade (Figure S2b) revealed that the DDD motif is conserved within this clade. Several Angiosperms contain two or more AGOs with the DDD triad (Figure 1c and S1a). Hence, the duplication of DDD-containing AGOs has occurred several independent times in evolution, suggesting important and perhaps specific functions in higher plants.

### AGO2 and AGO3 show cell-specific expression during reproduction

The *AGO2* and *AGO3* promoters do not display similarities despite the otherwise high conservation of their coding sequence (Figure 1a), suggesting distinct expression patterns. To address this question in detail, we stably expressed full-locus fluorescent protein fusions under native promoters (the promoter sequences used are depicted in Figure 1a), referred to as p*AGO3:mCherry-AGO3* and p*AGO2:mCherry-AGO2*. In light of online public expression profiles, we primarily focused our analyses on reproductive tissues (Winter *et al.*, 2007). The expression of the reporter constructs was imaged in plants co-expressing *LIG1-GFP* as a ubiquitous nuclear marker, in order to facilitate tissue recognition within the developing seeds.

p*AGO3:mCherry-AGO3* expression occurred in a few cells in the chalazal integument of ovules and seeds (Figure 2a-b), consistent with previously published laser-capture (LCM) expression data (Belmonte *et al.*, 2013). The fluorescent signal of p*AGO3:mCherry-AGO3* increases during early stages of seed development (Figure 2a-b). We could not detect mCherry-AGO3 signal within the female gametophyte, the male gametophyte, or in the embryo and endosperm, which develop after fertilization. p*AGO3:mCherry-AGO3* expression was detected in regions proximal to vasculature termination sites both at the end of stamen filaments (Figure 2c) and at the base of floral meristems (Figure 2d). In sharp contrast, p*AGO2:mCherry-AGO2* expression in analyzed reproductive organs was germline-specific, with high signals in egg cells of the female gametophyte (Figure 2e) and in sperm cells of the pollen grain before fertilization (Figure 2f). Note that the foci observed in the sperm cells may result from previously documented aggregation artifacts caused by mCherry or cytoplasmic concentration rather than *bona fide* cytoplasmic structures or compartments (Katayama *et al.*, 2008; Kremers *et al.*, 2011). Following fertilization, we could not detect p*AGO2:mCherry-AGO2* signal in the developing endosperm nor in the developing embryo.

**Figure 2.**
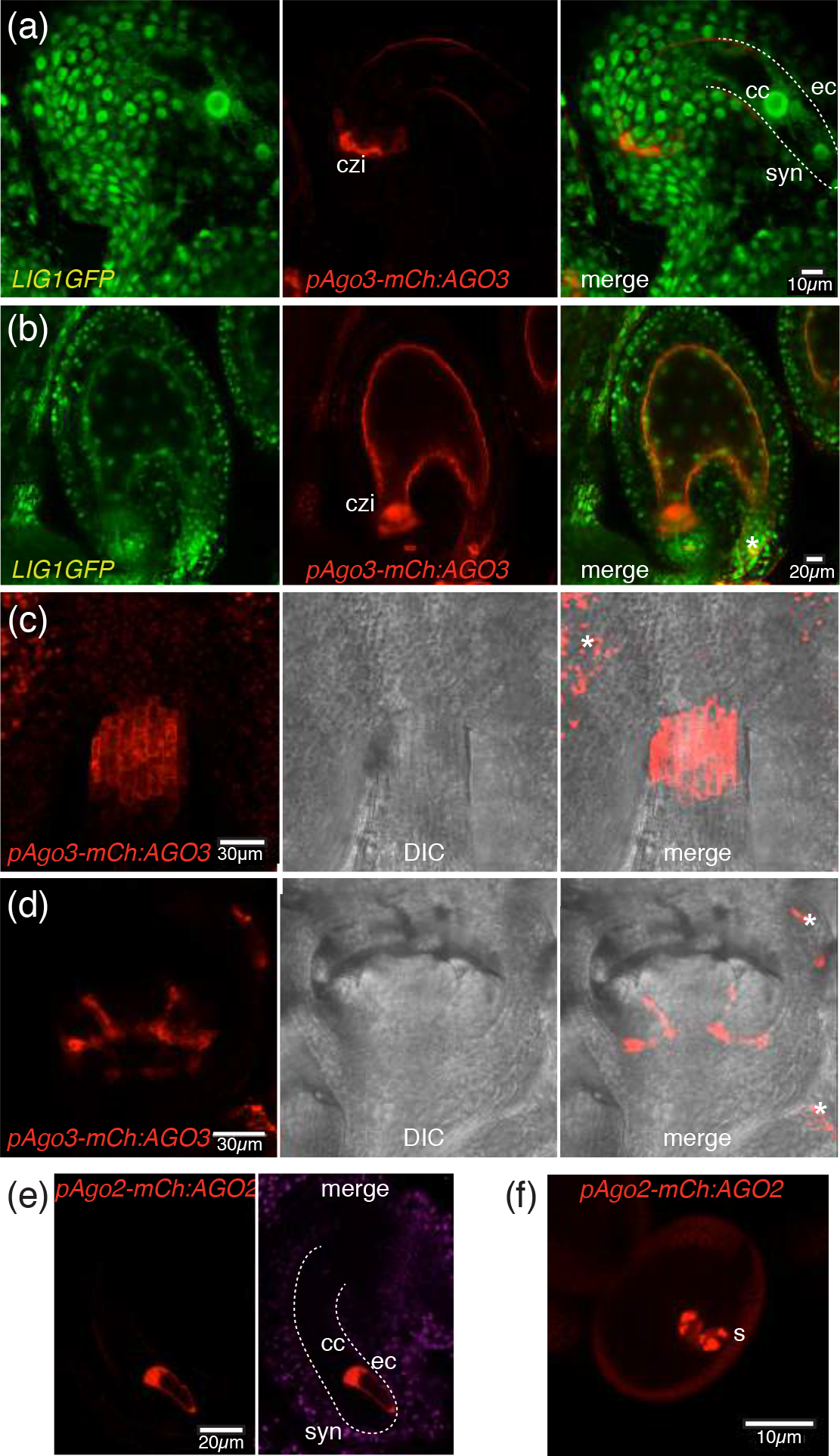
AGO3 and AGO2 display cell specific expression in reproductive tissues. (a-f) Confocal images from transgenic plants expressing a *pAGO3:mCherry-AGO3* (a-d) and *pAGO2:mCherry-AGO2* (e-f). mCherry-AGO3 is expressed in the chalazal integument in ovules (a) and seeds (b); at the end of stamen filaments (c) and at the base of the floral meristem (d). mCherry-AGO2 is expressed in the egg cell (e) and sperm cells(f). Scale bars are shown as a white rectangles. czi-chalazal integument, s-sperm cells, ec-egg cell, cc-central cell; syn-synergids; asterisks correspond to autofluorescence; DIC-Differential-Interference-Contrast.

The *AGO2* mRNA is a known target of miR403 (Allen *et al.*, 2005). The *AGO3* transcript also contains a putative miR403 binding site located in the 3’UTR (Figure S3). To test if and how miR403 influences *AGO2* and *AGO3* expression patterns, we engineered plants expressing transcriptional reporters for *AGO2* and *AGO3*, respectively named p*AGO2:H2B-mCherry* and p*AGO3:H2B-mCherry*. Both transcriptional reporters contained promoter sequences up to the translation start site, thus including the 5’UTR, but excluding the 3’UTR and were, therefore, devoid of the miR403 target site. Both reporters showed similar expression patterns in reproductive tissues compared to the respective full genomic fluorescent construct (Figure 3a-b). This result therefore excludes a strong miR403 contribution to the tissue-specific expression of AGO2 and AGO3. miR403 most likely regulates *AGO2* and *AGO3* expression level in their cognate tissues of expression. Consistent with this interpretation, both *AGO2* and *AGO3* transcripts are up-regulated in miRNA-deficient mutants of *Arabidopsis* including *ago1-27, hen1-6* and *dcl1-11* (Figure 3c-d). Furthermore, as the expression pattern between transcriptional and translational fusions are similar and that neither AGO2 nor AGO3 is detected outside its cognate expression domain, it strongly suggests that both proteins are cell-autonomous, at least in the tissues inspected.

**Figure 3.**
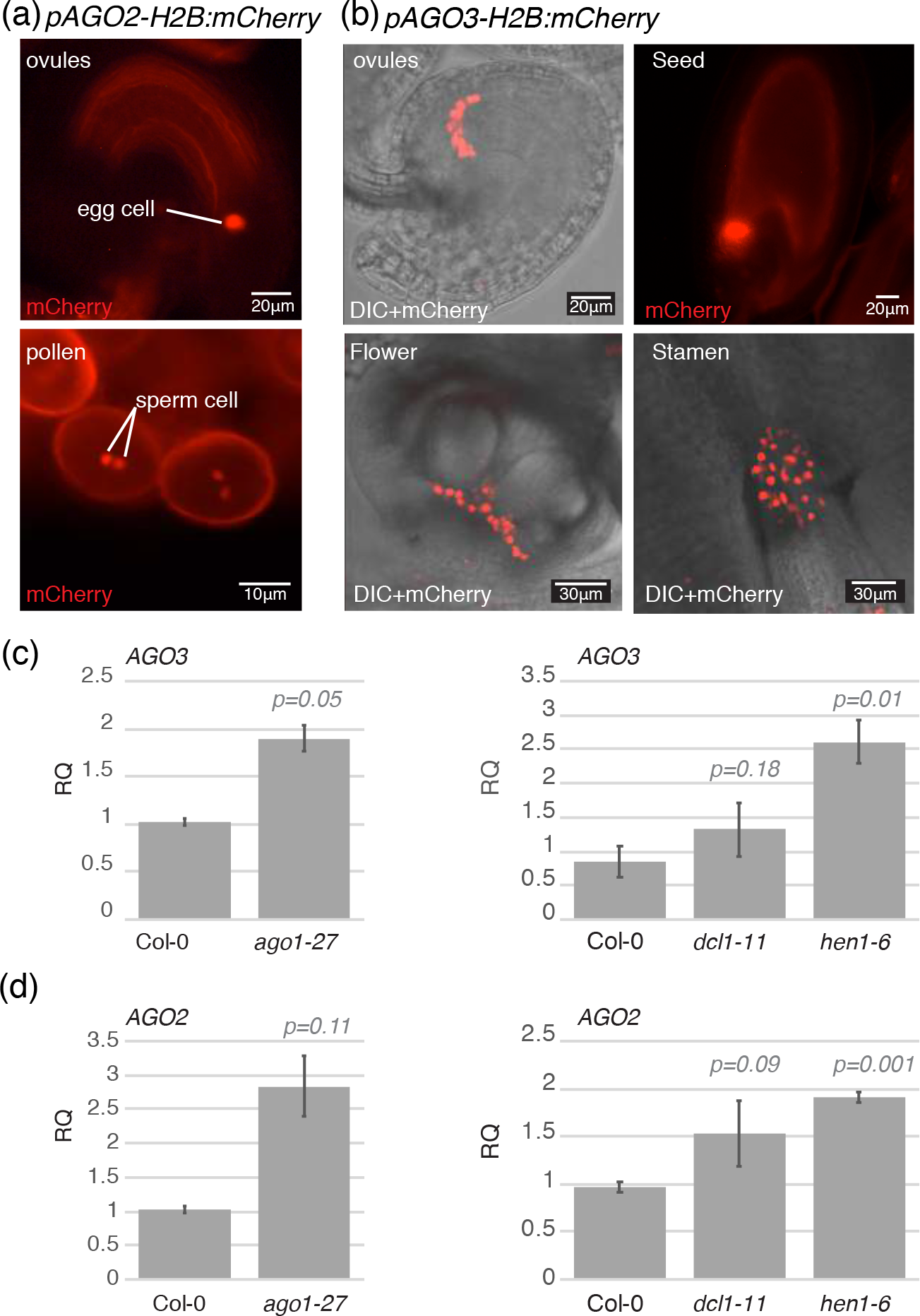
*AGO2* and *AGO3* regulation by mir403 affect expression level but not pattern. (a) Epifluorescence images showing *pAGO2:H2B-mCherry* expression in the egg cell and sperm cells. (b) Confocal images showing *pAGO3:H2B-mCherry* expression in the chalazal integument of the ovule. *pAGO3:H2B-mCherry* is also expressed at the base of flower primordia and end of stamen filament. Epifluorescence images showing *pAGO3:H2B-mCherry* expression in seed. (c) qPCR showing upregulation of *AGO3* in *ago1-27, dcl1-11* and *hen1-6* mutant in inflorescences. (d) qPCR showing upregulation of *AGO2* in *ago1-27, dcl1-11* and *hen1-6* mutant inflorescences. (c-d) Error bar represents standard deviation of two or three biological replicates. *ACT11/GAPC* were used as normalizer for qPCR. *p* indicates the *p* value obtained after a Student’s t-test compared to Col-0.

We conclude that AGO2 and AGO3 display non-overlapping expression patterns in reproductive tissues. AGO3 is expressed during early stages of seed development, in a specific subset of sporophytic cells located in the proximity of aerial vasculature terminations. AGO2, by contrast, is expressed in both male and female gametes.

### AGO3 binds 5’ Adenosine sRNA of both 24nt and 21nt

Sequencing of sRNAs from immunoprecipitates (IPs) has shown that AGO2 preferentially binds 21nt sRNAs with a 5’ Adenosine (Mi *et al.*, 2008). Upon salt stress, AGO3 was shown to bind 24nt with a 5’Adenosine bias. In order to investigate if AGO3 sRNA loading in silique was comparable to its loading upon salt stress, we generated stable *Arabidopsis* lines harboring p*AGO3:FHA-AGO3* construct, which expresses an N-terminal Flag-epitope-tagged version of AGO3 under its cognate promoter.

To first evaluate if FHA-AGO3 could indeed bind sRNAs *in planta* and to assess AGO3 binding affinity, we performed a transient infiltration in *N. benthamiana* of FHA-AGO3 together with a *p35S-GFP* construct and subsequent Flag-immunoprecipitation (Figure 4a). Infiltration of *p*35S-*GFP* results in the production of GFP sRNAs that can be analyzed by northern blotting. Both 21nt and 24nt GFP-derived sRNAs could be detected following AGO3 IP in opposition to only 21nt GFP-derived sRNAs following an AGO2 IP. AGO1 and AGO4 control IPs bound the expected sRNA sizes of 21nt and 24nt respectively. Our result shows that unlike AGO2, AGO3 can bind both 21nt and 24nt *in planta*.

**Figure 4.**
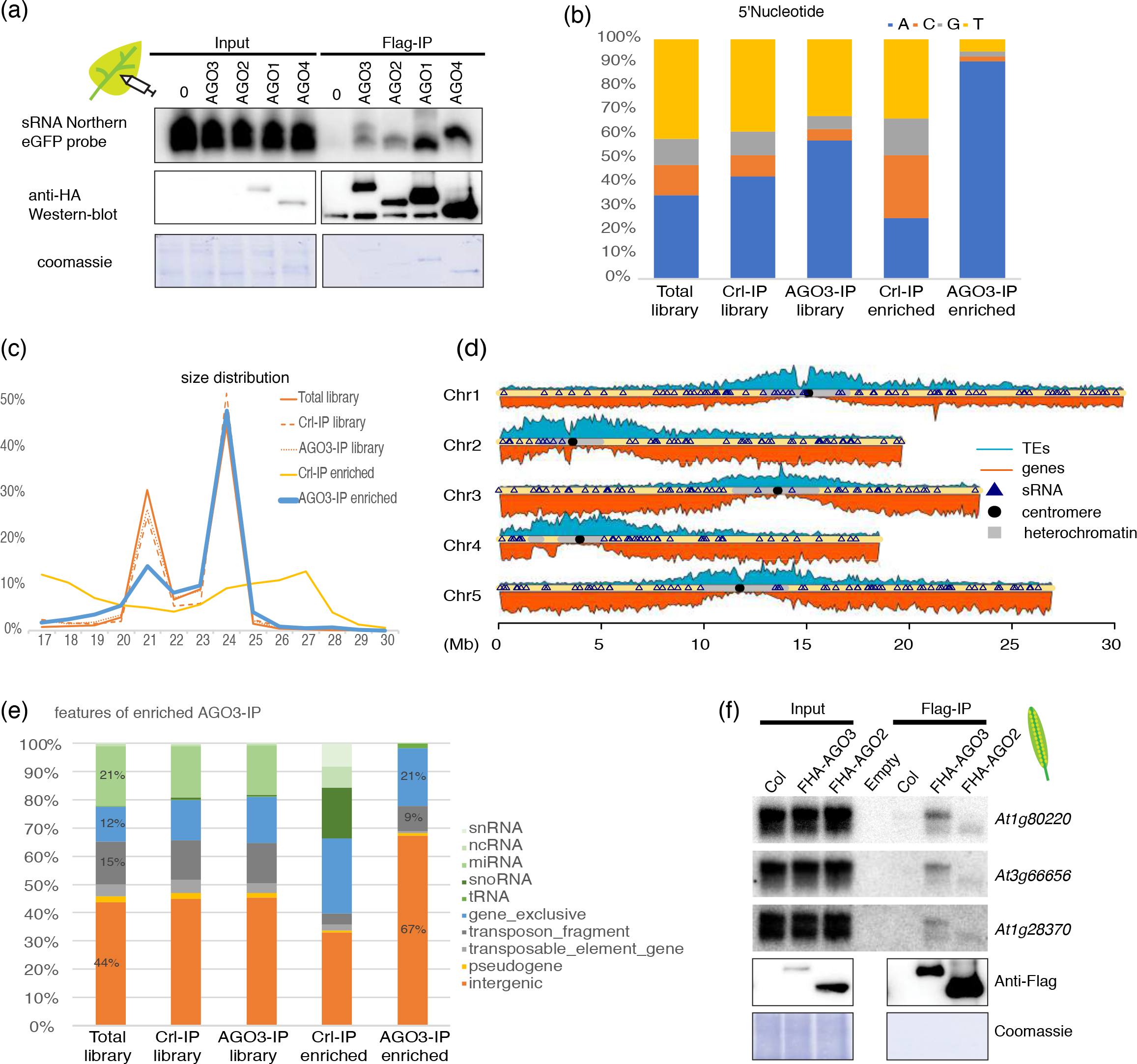
Characterization of AGO3-bound small RNA. (a) Northern blot showing that FHA-AGO3 can bind both 21nt and 24 nt sRNAs in transient expression in N. benthamiana. (b-e), AGO3-bound sRNAs were obtained by deep sequencing of a Flag IP on 1-5 DAP silique samples of *ago3-3 pAGO3:FHA-AGO3* transgenic plants. (b) AGO3 binds preferentially sRNAs with 5′A. (c) AGO3 binds 21- and 24-nucleotide sRNAs. (d) Chromosomic distribution of AGO3 bound sRNA. (e) Functional annotation of AGO3 bound sRNA. (f) Northern blot analysis of sRNAs after Flag IP in 4-6 DAP siliques of Col-0, *ago3-3 pAGO3:FHA-AGO3* and *ago2-1 pAGO2:FHA-AGO2* compared with input control.

In order to analyze FHA-AGO3 binding affinity in *Arabidopsis*, the p*AGO3:FHA-AGO3* construct was transformed in *ago3-3* mutant plants. sRNAs were isolated from Flag IPs conducted on 1-5 days after pollination (DAP) siliques of p*AGO3:FHA-AGO3* plants (referred to as AGO3 IP). sRNAs from a Flag IP in non-transgenic (Col-0) 1 −5DAP siliques (referred to as Ctrl-IP) and total RNA from (Col-0) 1-5 DAP siliques (referred to as Total) were used as control. sRNAs were subjected to Illumina sRNA deep sequencing (Figure 4 and S4). Because AGO3 expression is restricted to chalazal integuments in siliques, we anticipated a low signal-to-noise ratio due to inherent dilution effects. The sRNA contents of the libraries were aligned to the *Arabidopsis* genome and specific enrichment was calculated along 500bp genomic windows; sRNAs over-represented (10-fold enrichment) in the AGO3 IP compared to the Total were subsequently deemed as “AGO3-IP enriched”.

As a control, we similarly calculated a “Ctrl-IP -IP enriched” comparing the Ctrl-IP versus the Total. We found that, similarly to AGO2 IPs and AGO3 IPs upon salt-stress (Mi *et al.*, 2008; Zhang *et al.*, 2016), AGO3 exhibits a clear 5’ adenosine loading bias, since this property was shared by 90% of the sRNAs in the AGO3 enriched population (Figure 4b). However, unlike AGO2 (Mi *et al.*, 2008) and AGO3 during salt stress (Zhang *et al.*, 2016), AGO3 in siliques binds 24nt sRNAs as well as 21nt sRNAs (Figure 4c). Both, the 21nt and 24nt small RNAs enriched in the AGO3-IP display a 5’adenosine bias compared to the Total library and the control-IP enriched (Figure S5). As AGO3 binds a larger proportion of 24nt sRNAs compared to 21nt sRNAs, we analyzed the genome wide distribution of the AGO3-enriched fraction (Figure 4d) but could not detected any significant enrichment in heterochromatic regions but rather a uniform distribution along the five *Arabidopsis* chromosomes. The AGO3-enriched fraction revealed a marked depletion in miRNA compared to the Total sRNA fraction (Figure 4e). Unlike salt stress-bound AGO3 sRNA, there was no general enrichment in TE-derived sRNAs in the AGO3 enriched pool when compared to total sRNAs in siliques. In contrast, the AGO3-bound fraction was significantly enriched in sRNAs mapping to genes and intergenic regions (Figure 4e), which was confirmed directly for some loci by northern analysis of sRNAs extracted from Flag-AGO3 IP in 4-6 DAP siliques (Figure 4f). Northern results were consistent with the deep-sequencing data, including the relative 21nt/24nt abundance in AGO3-bound sRNAs (Figure S6). As a control, AGO2 IP did not bind 24nt sRNAs but only 21nt. Thus, AGO3 binds both 21nt and 24nt sRNAs in a ratio reflecting the accumulation of these sRNA species at their locus-of-origin. Furthermore, these 21nt and 24nt sRNA populations were overlapping, rather than separately distributed, in specific regions of the respective loci, suggesting their processing from a common double stranded RNA (dsRNA) precursor (Figure S6). Together, these results show that AGO3 binds sRNAs of 21nt and 24nt in siliques, which arise from genes and intergenic regions, and exhibit a strong enrichment for 5’ adenosine.

### AGO3 regulates gene expression in siliques

As AGO3 binds 21/24 nt sRNAs arising from genes in siliques, we wanted to investigate the effect of *ago3* loss-of-function on the transcriptome of *Arabidopsis* siliques. Two *ago3* mutant alleles have been described: *ago3-1*, with a T-DNA insertion in the last exon and the misexpression-allele *ago3-2* (Takeda *et al.*, 2008). We characterized a new mutant allele (GABI_743B03), named *ago3-3*, in which the *AGO3* transcript levels were at or below qPCR detection limit in all tissues inspected (cLv, inflo, 1-4 DAP and 5-8 DAP; Figure. 5a). To investigate AGO3 expression at the protein level, we raised a polyclonal antibody against immunogenic epitopes of the native protein. The AGO3 full-length protein has a predicted molecular mass of ~130kDa, and western blot analysis using an antibody against native AGO3 revealed a band at the expected size in wild-type plants, which was absent in all *ago3-3* mutant tissues analyzed, confirming the antibody’s specificity and knockout status of the *ago3-3* allele (Figure 5b). Moreover, the relative levels of AGO3 protein accumulation in wild-type plants were in agreement with the results of the qPCR and microscopy analyses, confirming that AGO3 is low in inflorescences and progressively up-regulated during early seed development within the siliques (Figure 5b). Noteworthy, the pattern of AGO3 accumulation in *ago2-1* remained globally unchanged (Figure 5b). Similarly, to AGO3 protein, AGO2 protein was neither up or down regulated in *ago3-3* mutant (Figure S7). Suggesting that AGO3 and AGO2 proteins are not subjected to cross-regulation or compensatory expression mechanisms.

**Figure 5.**
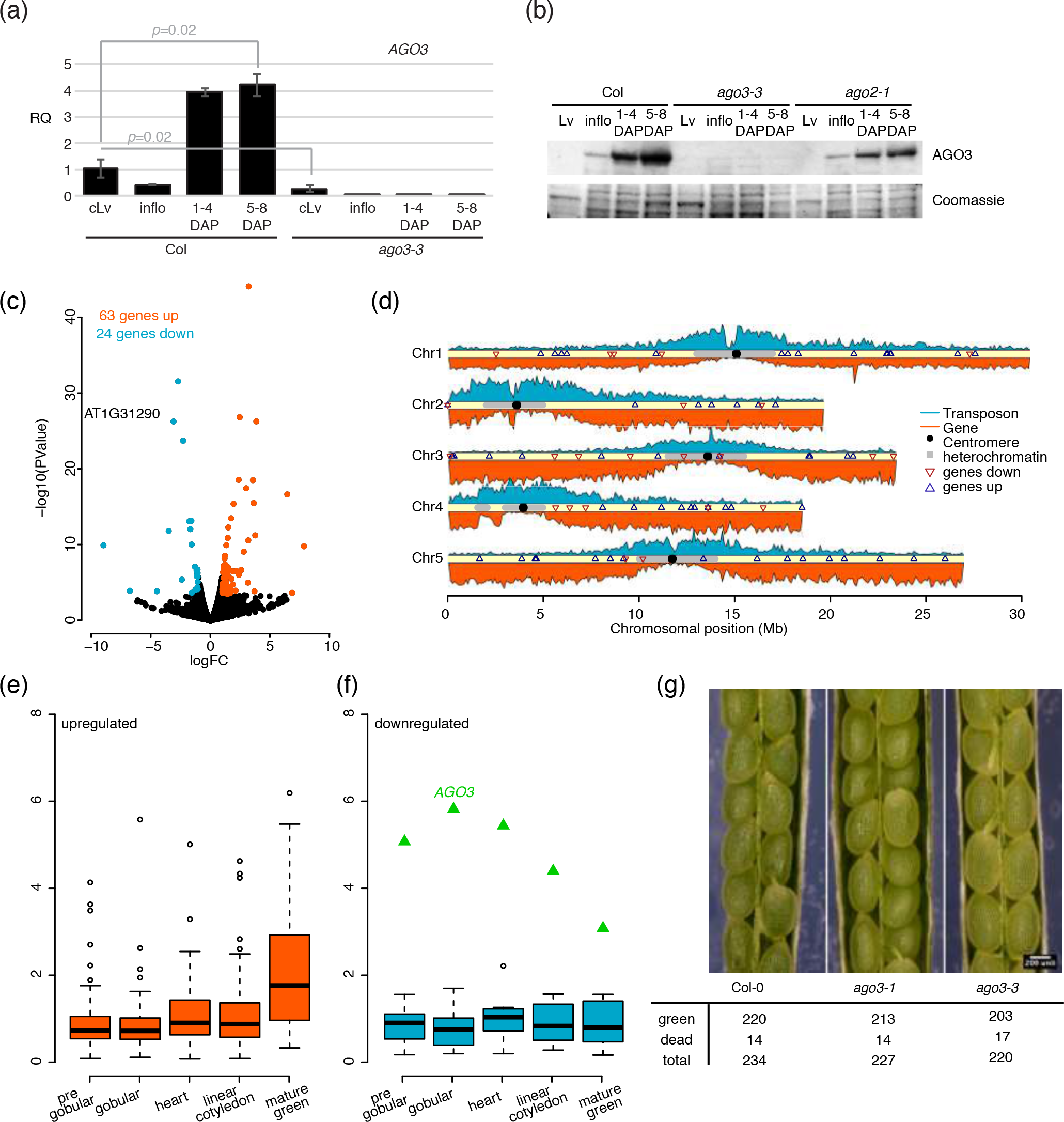
Transcriptome analysis of *ago3-3* mutant. (a) q-PCR analysis of *AGO3* expression in different tissues. Error bars represents standard error of two or three biological replicates. *p* indicates the *p* value obtained from a Student’s t-test. (b) Western blot analysis of AGO3 protein accumulation in different tissues and indicated genotype. (c) volcano plot showing the 87 mis-expressed genes in *ago3-3* compared to Col-0 1-5DAP siliques. (d) genomic location of *ago3-3* mis-expressed genes (down regulated genes are represented by red triangle and up regulated genes are represented by blue triangle). (e-f) Expression time course of up-regulated (e) and down (f) regulated genes in the chalazal seed coat at different stages of seed development. (g) Pictures and quantification of seed abortion at the green seed stage of Col-0, *ago3-3* and *ago3-1*. Lv, leaves; cLv, cauline leaves; Inflo, Inflorescence; DAP, day after pollination

To gain insight into potential roles for AGO3, we performed comparative RNA sequencing analyses on 1-5 DAP siliques of Col-0 wild-type and mutant *ago3-3* plants. RNA-seq libraries corresponding to two biological replicates from each genotype were sequenced using the Illumina technology. The analysis showed a higher number of up-regulated compared to down-regulated genes in *ago3-3* mutants (63 versus 24; Figure 5c), as expected from an anticipated function of AGO3 in gene silencing. Further investigation of the loci showing at least a 2-fold expression change (FDR < 0.05; Figure 5d) showed that *AGO3*-regulated genes were distributed along the five chromosomes’ arms.

To further characterize the effect of the *ago3-3* mutation, we analyzed the expression of up- and down-regulated genes during seed development, specifically in chalazal seed coat of wild-type samples, using data acquired by Belmonte *et al* (Belmonte *et al.*, 2013). For genes up-regulated in the *ago3-3* background, we observed that expression tends to increase during seed development, particularly at later stages (Figure 5e). In contrast, the expression of down-regulated genes in *ago3-3* remains nearly constant during the same time-window (Figure 5f). Moreover, the increased gene expression during seed development is anti-correlated with *AGO3* expression in the chalazal seed coat (Figure 5f), which decreases at later development stages. No such clear tendency was observed in the other seed compartments inspected. Variation in gene expression does not seems to result in developmental abnormalities, as no seed abortion was observed in the *ago3-3* mutant (Figure 5g).

In an attempt to identify direct targets of AGO3, we investigated potential overlaps between AGO3-bound sRNAs in siliques and the up-regulated genes found in the *ago3-3* mutant analysis (Figure S8). The only overlapping locus was *AGO3* itself, but as no sRNAs were found in the Col-0 Total library, the AGO3 sRNAs likely originate from the pAGO3:*FHA-AGO3* transgene. This lack of substantial overlap between the IP and transcriptome may have several causes: (i) the mRNA sequencing may have mainly identified indirect targets of *AGO3*, because the experiment was conducted on total siliques rather than isolated chalazal integument cells where *AGO3* is mostly, if not exclusively, expressed (Figure. 2), and/or (ii) sRNA-loaded AGO3 may act mainly as a translational repressor, in which case little or no impact on target mRNA accumulation would be expected.

### AGO3 is localized to the cytoplasm and co-sediments with polysomes

In order to get a better understanding of the molecular function of AGO3 in *Arabidopsis* cells, we investigated its intracellular localization. In *Arabidopsis*, AGO1 and AGO4 proteins are known to shuttle between the cytoplasm and the nucleus. However, their steady state localization seems to reflect their involvement in the TGS or PTGS pathways. Accordingly, AGO1 protein is mainly in the cytoplasm (Bologna *et al.*, 2018), whereas AGO4 mainly localizes to the nucleus (Ye *et al.*, 2012). We observed that the mCherry-AGO3 protein is mainly in the cytoplasm in both stamen filament and ovule integument cells (Figure 6).

**Figure 6.**
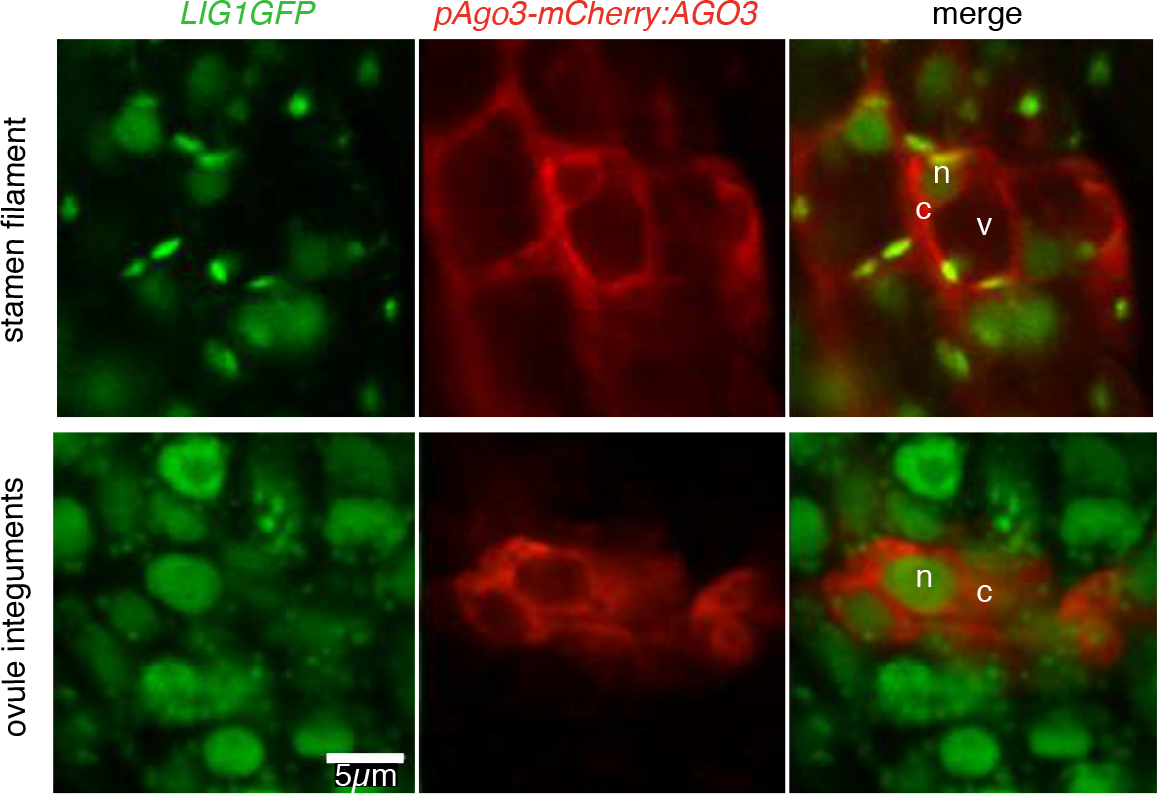
AGO3 localize to the cytoplasm. Confocal imaging showing *pAGO3:m-Cherry-AGO3* localisation in the cytoplasm of cells in the stamen filament as well as ovule integuments. *LIG1-GFP* is used as a DNA marker. n, nucleus; c, cytoplasm.

To get deeper insights into the possible AGO3 mode(s) of action, we performed mass-spectrometry analysis of AGO3 co-immunoprecipitated proteins. Flag IPs were conducted on 1-5 DAP siliques of *ago3-3* plants expressing p*AGO3:FHA-AGO3* in two biological replicates; Flag IPs conducted in parallel in non-transgenic Col-0 siliques provided the negative controls. Only proteins displaying a minimum of 2-fold enrichment in both biological replicates were selected, leading to a list of 79 AGO3 IP-enriched proteins that was noticeably devoid of any known member of the RdDM/TGS pathway (Table S1). In contrast, a GO-term enrichment analysis using agriGO (Du *et al.*, 2010) revealed a significant enrichment in NTP-dependent RNA helicases and core ribosomal constituents (Figure 7a). The latter finding prompted us to investigate whether AGO3 can associate with ribosomes, either as monosomes or polysomes (the latter reflecting active translation), where AGO3 could potentially achieve PTGS via translational repression as evoked in an earlier part of this study. Using conventional isolation procedures via differential centrifugation (Mustroph *et al.*, 2009), we found in 4-6 DAP siliques (Figure 7b) and in 1-5 DAP siliques (Figure S9a) that AGO3 co-sediments with both monosomes and polysomes as it has been shown for AGO1 (Figure S9a) (Lanet *et al.*, 2009). We then examined the presence of AGO3-bound sRNAs on the translation apparatus, using pools of the monosomes and polysomes fractions prepared above. Northern blotting revealed the presence of both the 21nt and 24nt forms of *AT1G80220*-derived sRNAs in polysomes (Figure S9b). These results suggest that AGO3 and AGO3-bound sRNAs specifically interact with the active translational machinery in siliques and could thus possibly regulate gene expression by PTGS *via* translational repression.

**Figure 7.**
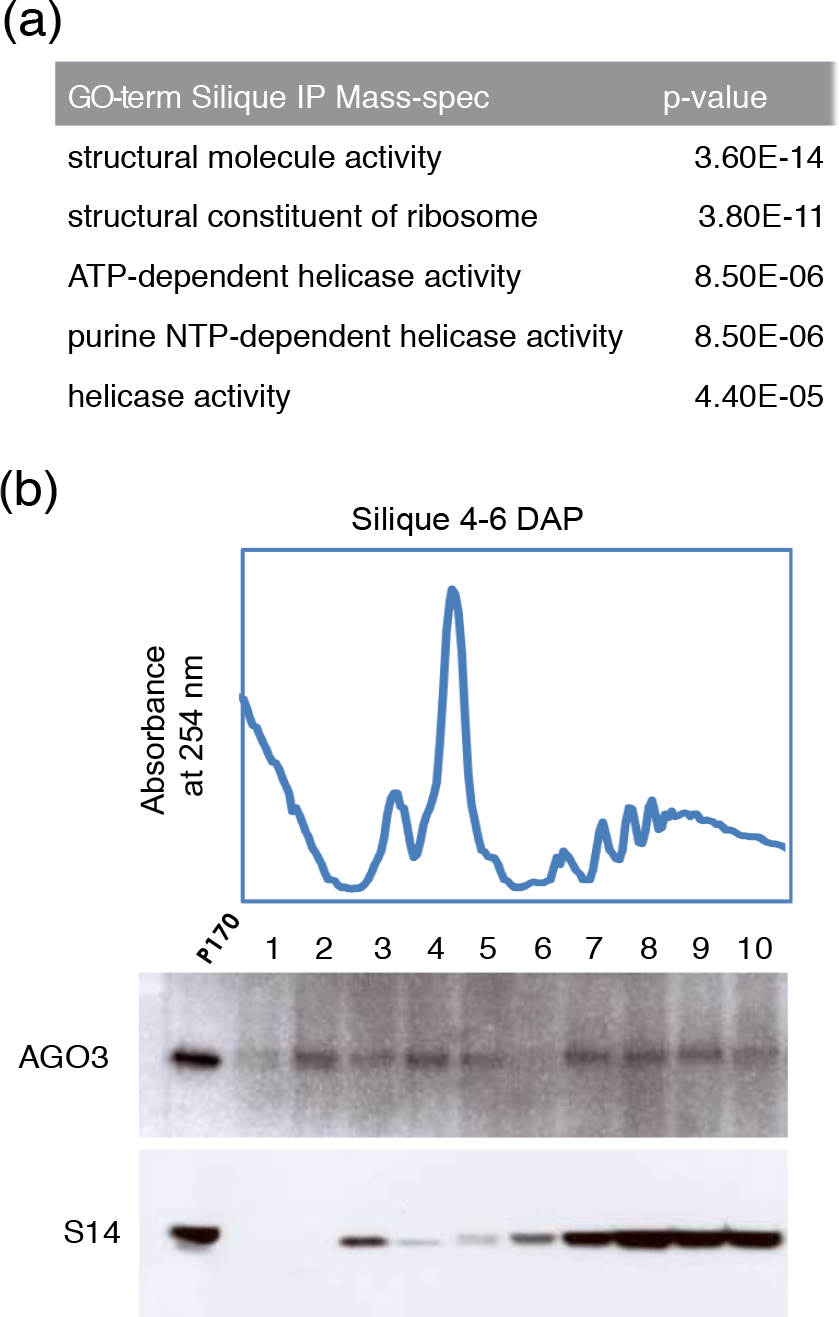
AGO3 co-sediments with the translation machinery. (a) Go-term enrichment of putative AGO3 interactors identified by mass spectrometry analysis of *ago3-3 pAGO3:FHA-AGO3* Flag IP in 1-5 DAP siliques. (b) Western blot analysis of polysome fractionation on 4-6 DAP siliques showing the co-sedimentation of AGO3 with monosomes and polysomes.

## DISCUSSION

Unlike for other *Arabidopsis* AGO proteins, the expression pattern, putative functions, and modes of action of AGO3 in native condition had remained mostly uninvestigated so far. Here, we discuss our findings in the context of AGO3’s closest and relatively well-characterized homolog, AGO2. *AGO3* arose from a recent transposon-driven duplication at the *AGO2* locus, an event restricted to the *Arabidopsis* lineage. Despite their high aminoacid sequence identity, the expression patterns of each protein differ drastically, due to their unrelated promoter sequences. As such AGO3 and AGO2 could be an example of subfunctionalization. In reproductive organs, AGO3 is expressed in siliques, and more specifically within the chalazal integument of developing seeds but not in the endosperm or in the embryo. AGO3 is also expressed in other terminal vascular structures found at the bases of stamen and floral meristems. AGO2, however, is expressed in the male (sperm cells) and female (central cell) germlines. Unlike that of AGO2, whose basal accumulation is also detectable in vegetative tissues, AGO3 expression is at, or below detection levels in leaves. AGO3’s confinement to vascular structures in the apical growing tissues could suggest a subtle and/or highly specialized role in antiviral defense, since the phloem is the channel employed by most plant viruses for systemic infection and unloading into sink tissues.

AGO2 and AGO3 are representatives of a DDD motif-containing AGO clade, which seems to be conserved throughout higher plants. Here, we show that similarly to AGO2, AGO3 binds sRNAs with a strong 5’ nucleotide preference towards adenosine (~90% of the cases). One outstanding property of AGO3, not displayed by AGO2, is its ability to bind both 21nt and 24nt sRNAs produced from genes and intergenic regions. This binding reflects the accumulation of the respective sRNA species at their locus-of-origin. Furthermore, many sRNA types bound by AGO2, including tasiRNAs, miRNAs, and miRNA*s (Mi *et al.*, 2008) are not overrepresented in the AGO3 enriched fraction. Given their highly specific and non-overlapping expression patterns, this difference in sRNAs identity loaded into AGO2 and AGO3 may reflect distinct sRNA compositions in the cognate expression domains of each protein as well as different loading properties.

Unlike AGO3 loaded sRNAs upon salt-stress, the silique AGO3-bound sRNAs noticeably lack an enrichment for TE-derived 24nt sRNAs. Mis-expression of AGO3 in an *ago4-1* mutant was shown to partially complement the *ago4-1* DNA methylation defect, supporting a role of AGO3 in TGS (Zhang *et al.*, 2016). Directly testing the AGO3 effect on DNA methylation in siliques would be confounded by its highly cell-specific and discrete expression pattern but the hypothesis that AGO3 may mediate TGS cannot be excluded. A role for AGO3 in PTGS is supported by the fact that AGO3 belongs to the same clade as AGO2 and AGO7, two PTGS effectors that, like AGO3, exhibit slicing activities *in vitro* when loaded with exogenous 21-22nt siRNAs (Schuck *et al.*, 2013). AGO2 and AGO7 slicer-deficient transgenes do not complement the viral hypersusceptibility of the *ago2* mutant or the *ago7* mutant’s developmental defects, respectively, showing that slicing is required for their proper physiological function. The catalytic residues of AGO3 are fully conserved, suggesting that it can also mediate PTGS via slicing *in planta*. It will be interesting to test whether AGO3’s loading with 24nt siRNAs also allows slicing *in vitro*. Furthermore, AGO3 cytoplasmic localization and the absence of any RdDM component in the mass spectrometry analysis do not support a role for AGO3 in mediating RdDM, at least under normal growth conditions in siliques.

We have pointed out the poor overlap between AGO3-bound sRNAs in siliques and the list of up-regulated mRNAs found in the same tissues of *ago3-3* mutant plants. We believe there are at least two non-mutually exclusive explanations for this discrepancy: (i) a dilution effect due to the highly cell-specific expression of AGO3 in the chalazal integument, and/or (ii) a possible translational repression effect of AGO3 that would not be diagnosed by RNA sequencing. Of the two possibilities, the second is at least indirectly supported by our finding that AGO3 associates with components of the translation apparatus and indeed co-sediments with monosomes and polysomes, like the PTGS effector AGO1 (Figure 7; Table1). Moreover, both the 21nt and 24nt sRNA species derived from *AT1G80220* were found on polysomes. Interestingly, previous research revealed how an unconventional 24nt siRNA could guide translational repression of an HD-ZIP transcription factor mRNA in maize and a GFP reporter in Arabidopsis (Klein-Cosson *et al.*, 2015), further fueling the hypothesis that AGO3 may exert a function in PTGS. However, AGO3 has not been investigated in the latter study. Under normal growth conditions, AGO3 is expressed in a limited number of cells in the apical part of the plant. Interestingly, these cell types coincide with vasculature terminations, and, given the ability of endo-siRNA to move systemically (Molnar *et al.*, 2010), it is tempting to speculate that AGO3 could act as a filter, perhaps providing a safeguard against the entry of specific sRNA into gametic or embryonic tissue. Indeed, all nutrients unloaded from the phloem transit through the chalazal integument toward the endosperm to finally reach the embryo. Especially considering the sRNA binding competition that might exist between AGO3 and AGO4 (both loading 24nt with 5’A). However, our attempts to investigate this small RNA movement did not allow us to confirm this hypothesis. An alternative hypothesis could be that AGO3 act at vascular terminations to regulate gene expression that might be caused by phloem unloading of diverse other molecules than sRNAs, such as mobile mRNA or hormones.

## EXPERIMENTAL PROCEDURES

### Plant Materials and Growth Conditions

After three days at 4°C in the dark, seeds were germinated and grown on soil. Plants were grown under long days at 20-21°C (16h light/8h night). All plants were in Columbia (Col-0) accession. The mutants described in this work correspond to the following alleles: *ago1-27* (Morel *et al.*, 2002), *ago2-1* (Salk_037548), *ago3-2* (SALK_005335, (Takeda *et al.*, 2008)) *ago3-3* (GABI_743B03), *dcl1-11* (ZHANG *et al.*, 2008), *hen1-6* (SALK_090960, (Li *et al.*, 2005)). The insertion lines were provided by The Nottingham Arabidopsis Stock Centre (NASC) (http://arabidopsis.info/).

### Microscopy

Fluorescence images were acquired using laser scanning confocal microscopy (Zeiss LSM780) or Leica epifluorescence microscope. Brightness and contrast were adjusted using ImageJ (http://rsbweb.nih.gov/ij/) and assembled using ImageJ or Adobe Photoshop.

### Plasmid Construction and Transformation

All fragments were amplified by PCR using the Phusion High-Fidelity DNA Polymerase (Thermo). Primer sequences can be found in Supplementary Table 2. All plasmids were transformed into wild-type Columbia plants, *ago3-3, ago2-1* and/or LIG1-GFP marker line (Andreuzza *et al.*, 2010). All constructs were generated using Multisite Gateway technology (Invitrogen). *A. thaliana* transformation was carried out by the floral dip method (Clough and Bent, 1998). At least ten transgenic lines were analyzed, which showed a consistent fluorescence using a Leica fluorescent microscope or consistent level of FlagHA by western blot. Three independent lines with single insertions, determined by segregation upon BASTA selection, were used for further detailed analysis.

### RNA and qPCR analysis

Total RNA was extracted either with Qiagen RNeasy mini kit for silique samples or TRIzol reagent (Invitrogen) for other tissues. Total RNAs were DNase treated (DNaseI, Thermo Scientific) and reverse transcribed into cDNA using the Maxima First-Strand cDNA Synthesis kit (Thermo Scientific). Results were normalized to *GAPC* levels for seedlings, and to *GAPC* and *ACT11* for inflorescence and silique tissue. qPCR reactions were performed using KAPA fast Master Mix on a LightCycler480 II (Roche). Primers are listed in Table S2. The Relative Quantification value (RQ) represents the average RQ and the error bars represent standard error from at least two biological replicates. P values were calculated using a Student’s t-test. Total RNA was depleted of ribosomic RNA and libraries prepared and subjected to paired-end sequencing using the corresponding Illumina protocols at the Functional Genomics Center Zurich (http://www.fgcz.ch/). For sRNA deep-sequencing analysis, sRNAs were eluted from the Flag beads using TRIzol and precipitated with glycogen and isopropanol overnight at −20°C. Total sRNAs and IP sRNAs were processed into sequencing libraries and sequenced by Fasteris (http://www.fasteris.com, Switzerland).

### Protein and Immunoprecipitation analyses

Protein extraction and Western blot analysis were performed as previously described (Marí-Ordóñez *et al.*, 2013). Antibodies used in this paper are: Monoclonal ANTI-FLAG M2 Peroxidase HRP antibody (SIGMA A8592), Anti-HA-Peroxidase High Affinity 3F10 (ROCHE 12 013 819 001), S14 (Agrisera AS09 477), and Anti-AGO2 (Garcia *et al.*, 2012). For the native anti-AGO3, peptide antibodies were prepared in rabbits according to the DoubleX program of Eurogentec. Peptides used for AGO3 antibody production and immunization protocol were H-CRG FVQ DRD GGW VNP G-NH2 and H-CGH VRG RGT QLQ QPP P-NH2 both situated on the N-terminus of the AGO3 protein.

AGO3 protein immunoprecipitations were performed as previously described (Marí-Ordóñez *et al.*, 2013) with the following modifications: No preclearing was performed. Flag immunoprecipitation was performed using 30μl of EZview Red ANTI-FLAG M2 Affinity Gel from Sigma (SIGMA F2426) in 1.5ml lysate for 2-3h at 4C. Immunoprecipitation in non-transgenic plants or non-treated plants were used as a background control for all experiments. IP for the northern blot was conducted using antiFLAG-conjugated agarose beads (Sigma) and the supernatant was pre-cleared for 15 minutes with 50μl of Protein A-agarose beads (Sigma).

For Mass spectrometry analysis, protein complexes were washed once with IP buffer and twice with 1X TBS buffer and subsequently eluted from the beads using competition with FLAG peptide according to the manufacturer’s instructions (Sigma). The elution was precipitated, trypsin treated, and run using LC/ESI/MS/MS at the Functional Genomics Center Zurich. Mass spectrometry data analysis was performed using the Scaffold software (Proteome Software) with the following settings: Protein identification threshold of 5% FDR, minimum peptide 1, and peptide threshold 95%.

### RNA blot analysis

RNA from input and IP samples, suspended in 50% formamide, was separated on a 17.5% polyacrylamide-urea gel, electrotransfered to a HyBond-NX membrane (GE Healthcare), and crosslinked with 1-ethyl-3-(3-dimethylaminopropyl) carbodiimide-mediated chemical crosslinking, as previously described (Pall and Hamilton, 2008). Radiolabeled probes were made by incubating gel-isolated PCR fragments with the Prime-A-Gene kit (Promega) in presence of [α-^32^P]-dCTP (Hartmann Analytic). Multiple probes were tested on individual membranes by stripping with boiling 0.1% SDS and rehybridizing. Primers used to amplify the probes can be found in Supplementary Table 2.

### Sucrose density gradient fractionation of polysomes

Sucrose gradients were conducted according to the protocol of Mustroph *et al* (Mustroph *et al.*, 2009).

### Bioinformatics

#### sRNA-seq analysis

The Dot plot, the protein alignment, and the phylogenetic tree (neighbor joining method with 100 bootstraps) were generated using CLC genomic workbench 8. Putative AGO protein sequences were downloaded from Phytozome (http://phytozome.jgi.doe.gov) (Goodstein *et al.*, 2012) with a double Pfam domain filter (PIWI, PF02171 and PAZ, PF02170). Illustration for chromosomal loci positions, volcano plot and box plots were implemented with R and/or in-house scripts.

For sRNA sequencing analysis, the trimmed sRNA reads were aligned against chloroplast, mitochrondrial, rRNA, and tRNA sequences, and sequences of two chromosomal regions, which exhibit unusually high sRNA association (Chr2:1.10000 and Chr3:14194000.14204000) and which likely represent degradation products of spurious rDNA transcription. For alignment, bowtie (Langmead *et al.*, 2009) with the following parameters was used: −v 2–best −m 1000. The unaligned reads were kept for further processing. Subsequently, all the reads shorter than 17 nt and longer than 30 nt were discarded using awk command (see TableS3a task A). We then aligned the filtered sRNA-seq reads against the TAIR10 *Arabidopsis thaliana* genome using bowtie with the following parameters: −v 2–best −m 1000. TableS3b contains a summary of the read alignment scores. Using samtools sort and index (Li *et al.*, 2009), the resulting bam files were sorted and indexed. To determine the length distribution of entire libraries, bam files were converted to sam files (by samtools view) and the length distribution was extracted using a customized command line command based on the command line tool awk (see TableS3a task B).

Subsequently, the sum of sRNA-seq reads per 500 bp non-overlapping bin was assessed using HiCdat (Schmid *et al.*, 2015). For further analysis, genomic bins, which contained less than 5 reads in the total sRNA library were removed. The reads per bins values were normalized to the total read numbers across the libraries (using cpm() in the edgeR package (Robinson *et al.*, 2010)) To calculate the enrichment of AGO3 IPs, the number of reads per bin in the AGO3 IP was divided by the number of reads found in the total sRNA fraction. In parallel enrichment of the control FLAG IP over the total sRNA fraction was calculated. Later, significantly enriched bins had to fulfill following criteria: 10-fold enriched over the total sRNA fraction and less than 2-fold enrichment in the FLAG IP control fraction. For further analysis, sequences and coordinates of the significantly enriched bins were retrieved.

The first nucleotide of each alignment was obtained by customized awk script taking strand information of the alignment into account (for + alignments (see TableS3a task C); for – alignments see TableS3a task D). To resolve the first nucleotide identity by distinct sRNA sizes, above script was looped across sam files containing reads of a unique length only. To determine genomic features associated with the aligned sRNA reads, we extracted feature coordinates from the publicly available TAIR10 GFF3 file (TAIR10_GFF3_genes_transposons.gff, 49,811 kb, 2010-12-14). Using customized awk scripts, we added two features: promoters (500 bp up- and downstream of the start of the feature gene, respectively, depending on the genes orientation) (see TableS3a task E) and intergenic (all sequences that do not overlap the annotated GFF3 features and promoters), which were defined by using the bedtools complement tool (Quinlan and Hall, 2010) (complement of all features and entire chromosomes).

#### mRNA-seq analysis

For mRNA sequencing analysis, the filtered reads were aligned to the TAIR10 *Arabidopsis thaliana* genome using HISAT2 with default parameters (Kim *et al.*, 2015). After sorting and indexing using SAMTOOLS, the aligned RNA-seq reads were mapped to genomic features using the Rsubread (Liao *et al.*, 2013) package’s command RsubCounts() taking into account multi-mapping reads and strand specificity. The obtained gene read counts were subsequently analyzed for differential expression between the different genotypes (2 *ago3-3* homozygous, 2 *ago3-3* heterozygous, and 2 wild-type samples) using the edgeR (Robinson *et al.*, 2010) package. We only further analyzed genomic features, which exhibited at least 5 reads in at least 2 RNA-seq samples. The differential analysis was performed employing a linear model by using the estimateGLMCommonDisp() and estimateGLMTrendedDisp() functions. Differentially expressed features, which exhibited at least a 2-fold change and an FDR < 0.05 were scored as significantly differentially expressed.

For Figure 5e, microarray data from Belmonte *et al.* (Belmonte *et al.*, 2013) were extracted as pre-processed data from the Arabidopsis eFP Browser (Winter *et al.*, 2007). Expression means of the replicates were calculated for each transcript and represented as boxplots using R.

## ACCESSION NUMBERS

Data sets of small RNAs and RNA deep-sequencing generated in this study are deposited in the National Center for Biotechnology Information Gene Expression Omnibus (http://www.ncbi.nlm.nih.gov/geo/) under accession number XXXX.

## AUTHOR CONTRIBUTIONS

PEJ conceived the project. OV contributed to the experimental design. PEJ, NP, CC, CO and GS performed the research. SG and AM performed the bioinformatics analysis, together with PEJ. PEJ and OV wrote the manuscript with the help of NP.

## ACKNOWLEDGEMENTS

We thank the entire Voinnet lab for thoughtful discussions, constructive ideas and critical reading of the manuscript. We would like to thank the following people for their help: Andre Imboden for support concerning plant growth, Florian Brioudes and Christopher Brosnan for plasmids and advice during experiments. This project was supported by a core grant from ETH-Z. PEJ (Project 329404), NP (Project 299789) and CC (Project 275589) were supported by Marie Curie fellowship.

## SUPPORTING INFORMATION

Additional Supporting Information may be found in the online version of this article.

Figure S1. Additional phylograms

Figure S2. AGOs alignments highlighting catalytic residues.

Figure S3. miR403 binding details

Figure S4. Western blot of AGO3 IPs in 1-5 DAP siliques

Figures S5. 5’nucleotide bias according to small RNA size

Figures S6. Representation of the sRNA sequencing reads for the loci tested by Northern blot in Figure 4f

Figure S7. AGO2 expression in *ago3-3* mutant

Figure S8. Comparison between AGO3 enriched sRNA and *ago3-3* transcriptome

Figures S9. AGO3 and associated small RNA co-sediments with polysomes in 1-5 DAP siliques

Table S1. Raw output from AGO3 IP Mass Spectometry

Table S2. Oligonucleotides used in this study.

Table S3. Bioinformatic analysis information

---

**Figure S1.**
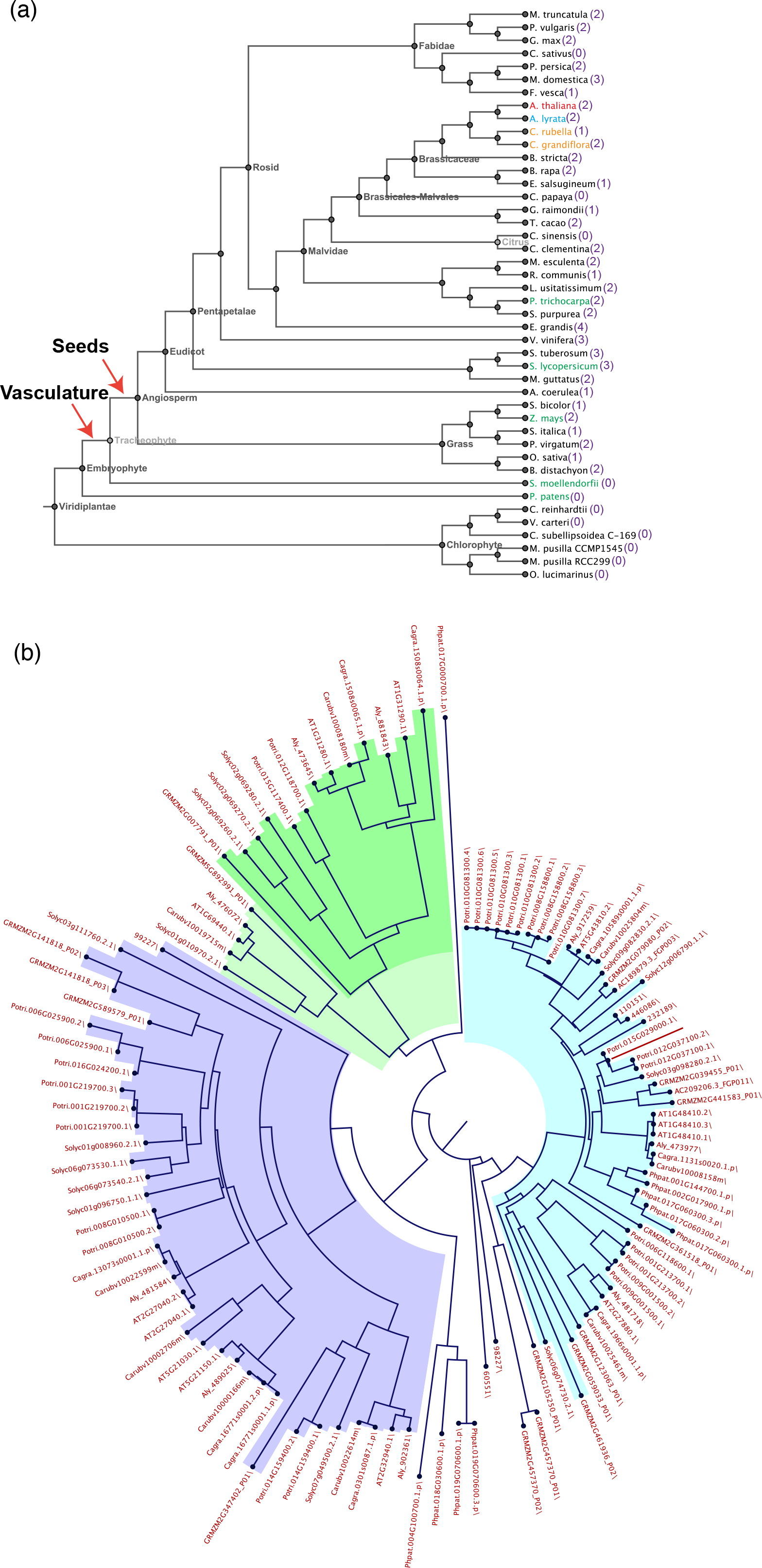
Additional phylograms. (a) Tree highlighting the species used to build the AGOs phylogenic tree. adapted from http://phytozome.jgi.doe.gov/pz/portal.html. A.thaliana is indicted in red, closely related species A.lyrata (blue) and Capsella spp. (C.rubella and C.grand flora in orange). Other representative species used are marked in green. AGO2/AGO3-like Argonautes containing a DDD motif are present in angiosperm but not in lower plants such as S.patens and S.moellendorf i. The number of DDD containing AGOs is indicated in purple. (b) Circular phylogram (Neighbor joining, bootstrap 100) showing that the duplication exists in Arabidopsis lineage but not in Capsella spp. Sequences used to build this tree were all obtained from http://phytozome.jgi.doe.gov/pz/portal.html and only putative proteins containing both PAZ and PIWI pfam domains were used. Classical Arabidopsis AGOs clades are represented: AGO2/3/7 in green, AGO4/6/8/9 in purple and AG01/5/10 in blue. Original protein sequence names from Phytozome are shown.

**Figure S2.**
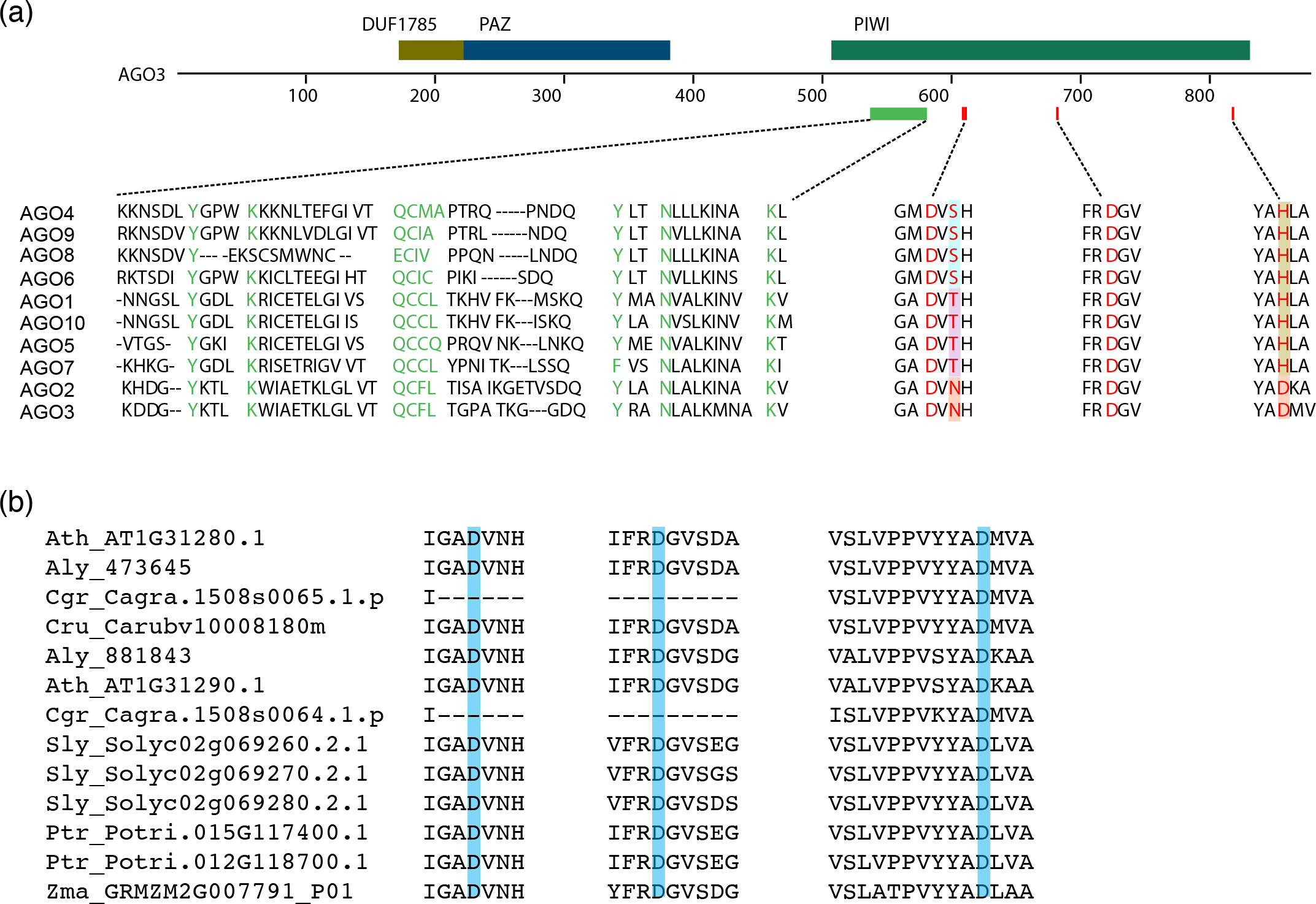
AGOs alignments highlighting catalytic residues. (a) Alignment of Arabidopsis Argonaute PIWI active residues (Red: catalytic residues, green: 5’ binding pocket residues). Highlighted residues show the unique DNDD motif of AGO3 and AGO2 as annotated on NCBI CDD cd02826 (http://www.ncbi.nlm.nih.gov/Structure/cdd/cddsrv.cgi?hslf=1&uid=cd02826). (b) Alignment of Argonaute PIWI active residues of the AGO2/AGO3 clade (Corresponding to the dark green clade in Figure 1 and Figure S1) highlighting their conserved DDD motif. Lack of some residues in C.grand flora is most likely due to incomplete genome annotation. Ath, Arabidopsis thaliana; Aly, Arabidopsis lyrata; Cgr, Capsella grandiflora; Cru, Capsella rubella; Sly, Solanum lycopersicum; Ptr, Populus trichocarpa; Zma, Zea mays.

**Figure S3.**
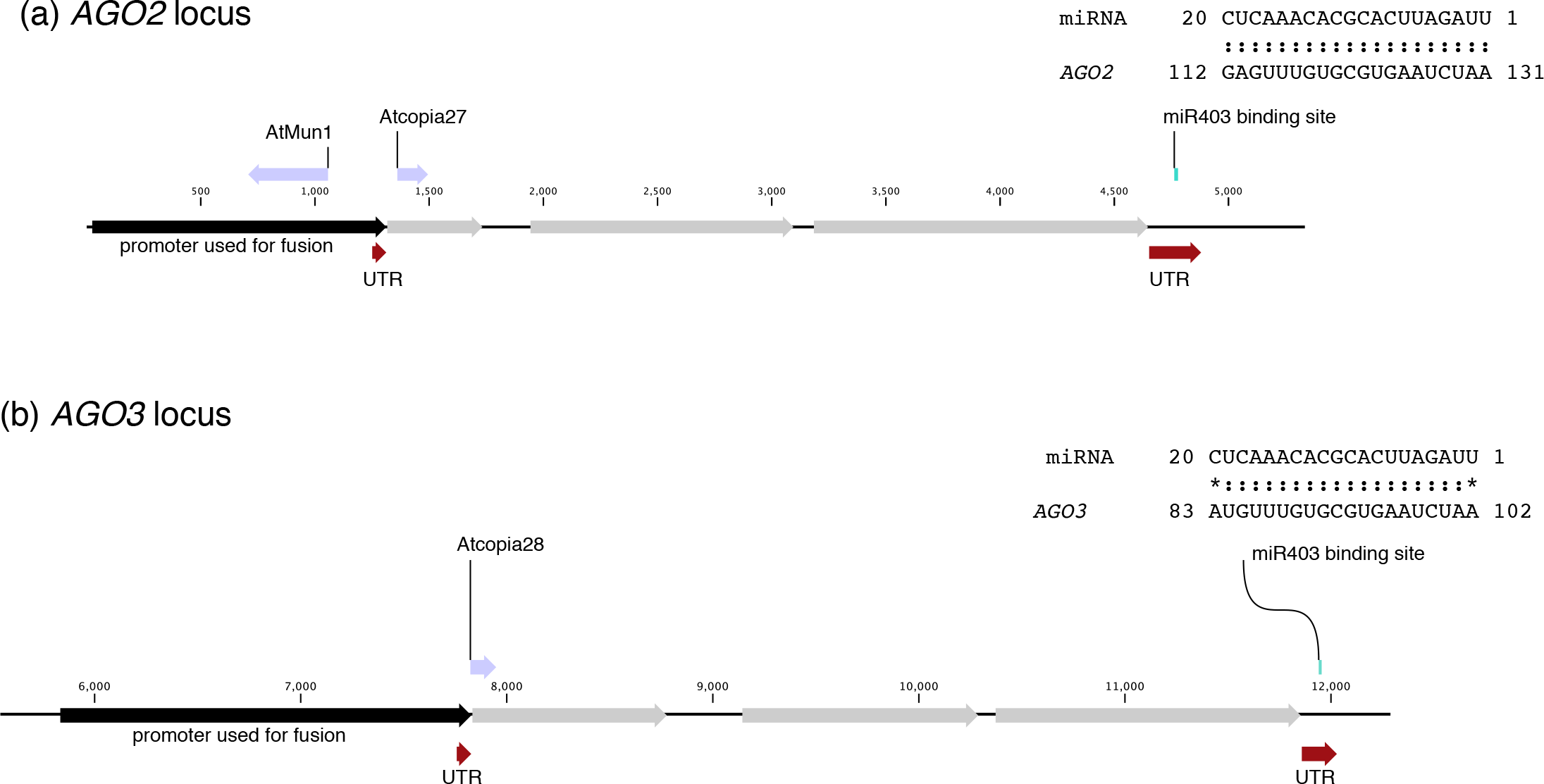
miR403 binding details. Localisation and sequence of the putative binding site of miR403 within AGO2 (a) and AGO3 (b) 3’UTR.

**Figure S4.**
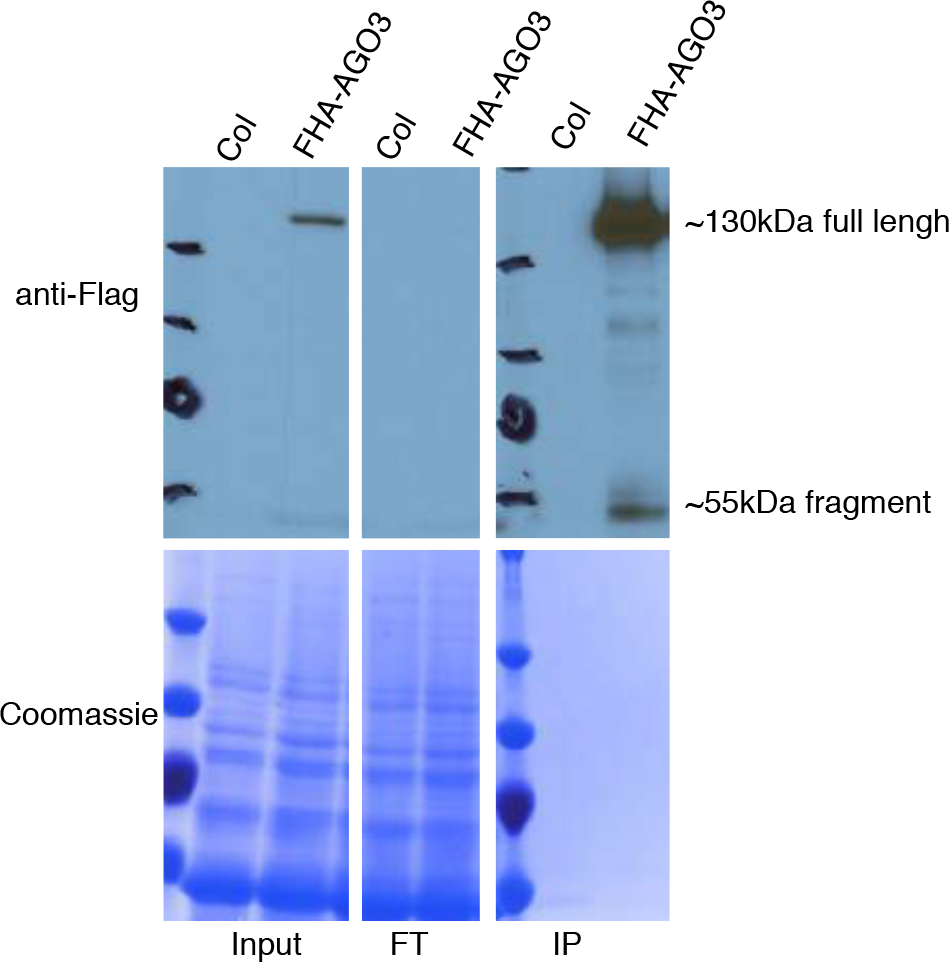
Western blot of AGO3 IPs in 1-5 DAP siliques. IP done in siliques from 1 to 5 DAP of ago3-3 pAGO3:FHA-AGO3 transgenic plants. Anti-Flag antibody was used to detect tagged AGO3 and Coomassie staining is used as loading control.

**Figure S5.**
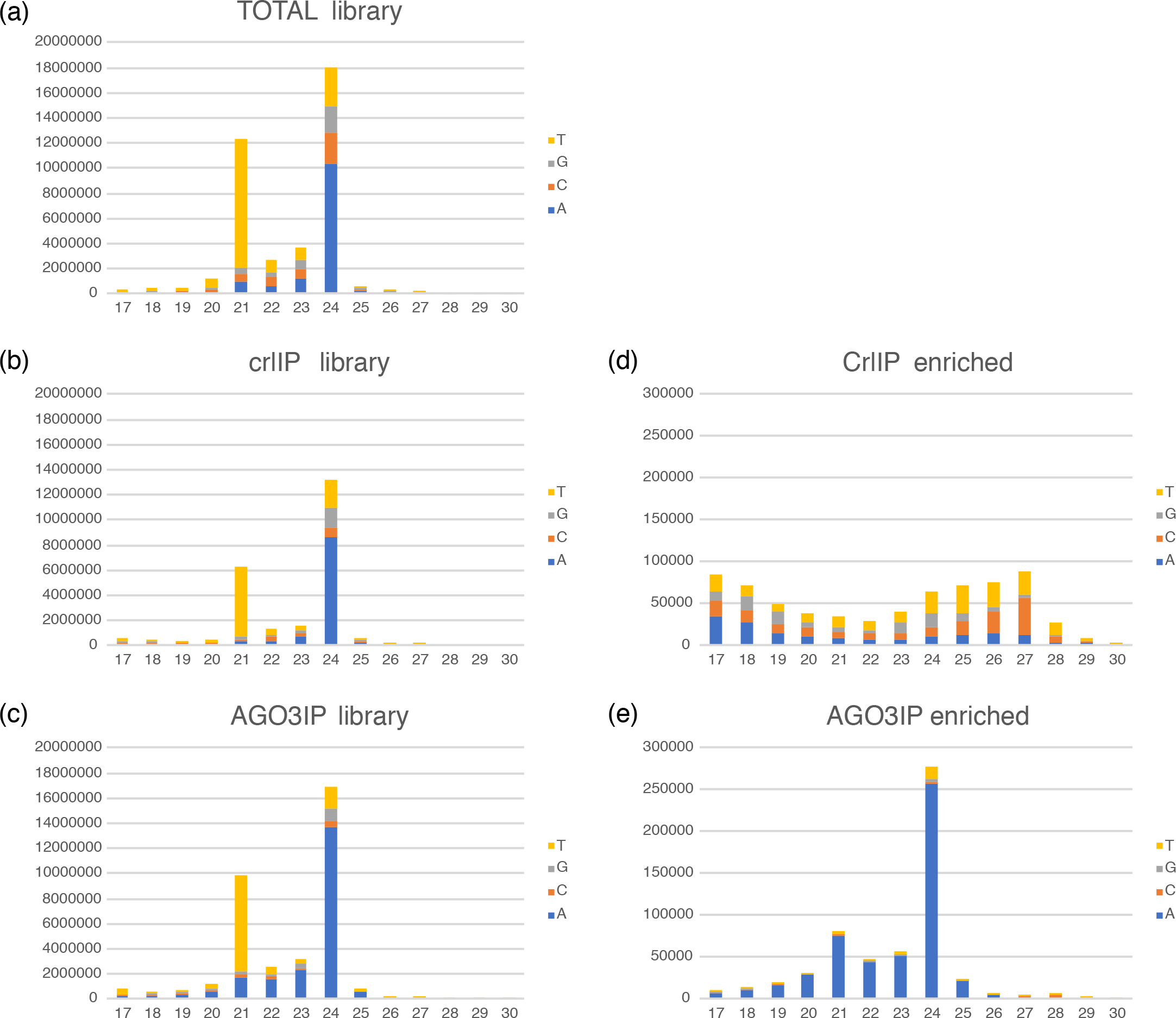
5’nucleotide bias according to small RNA size. 5’ nucleotide bias according to the small RNA size in the sequencing libraries as well as IP enriched fractions.

**Figure S6.**
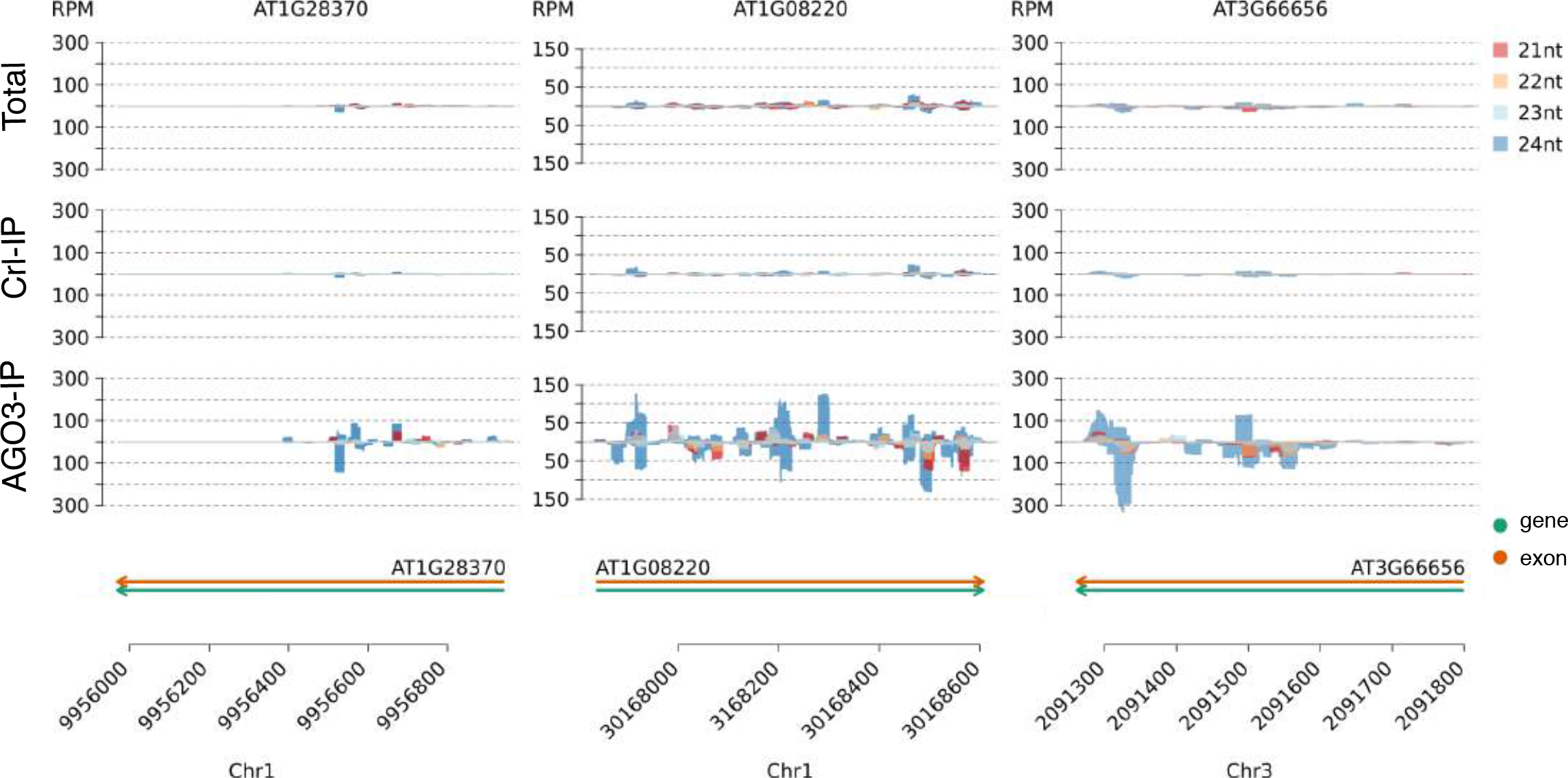
Representation of the sRNA sequencing reads for the loci tested by northern blot in Figure 4f. The color code indicates the size of the small RNA and the identity of the library is indicated on the left.

**Figure S7.**
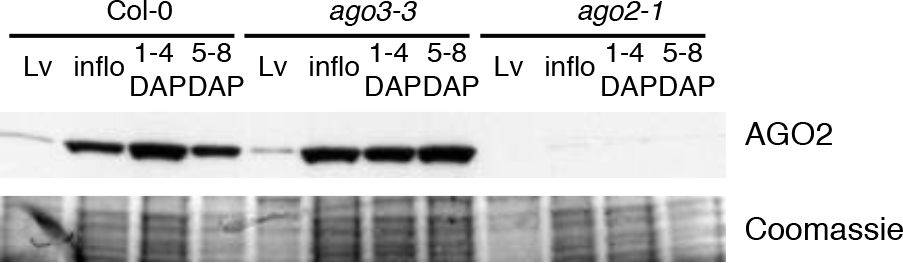
AGO2 expression in ago3-3 mutant. Western blot analysis of AGO2 protein accumulation in different tissues and indicated genotype.

**Figure S8.**
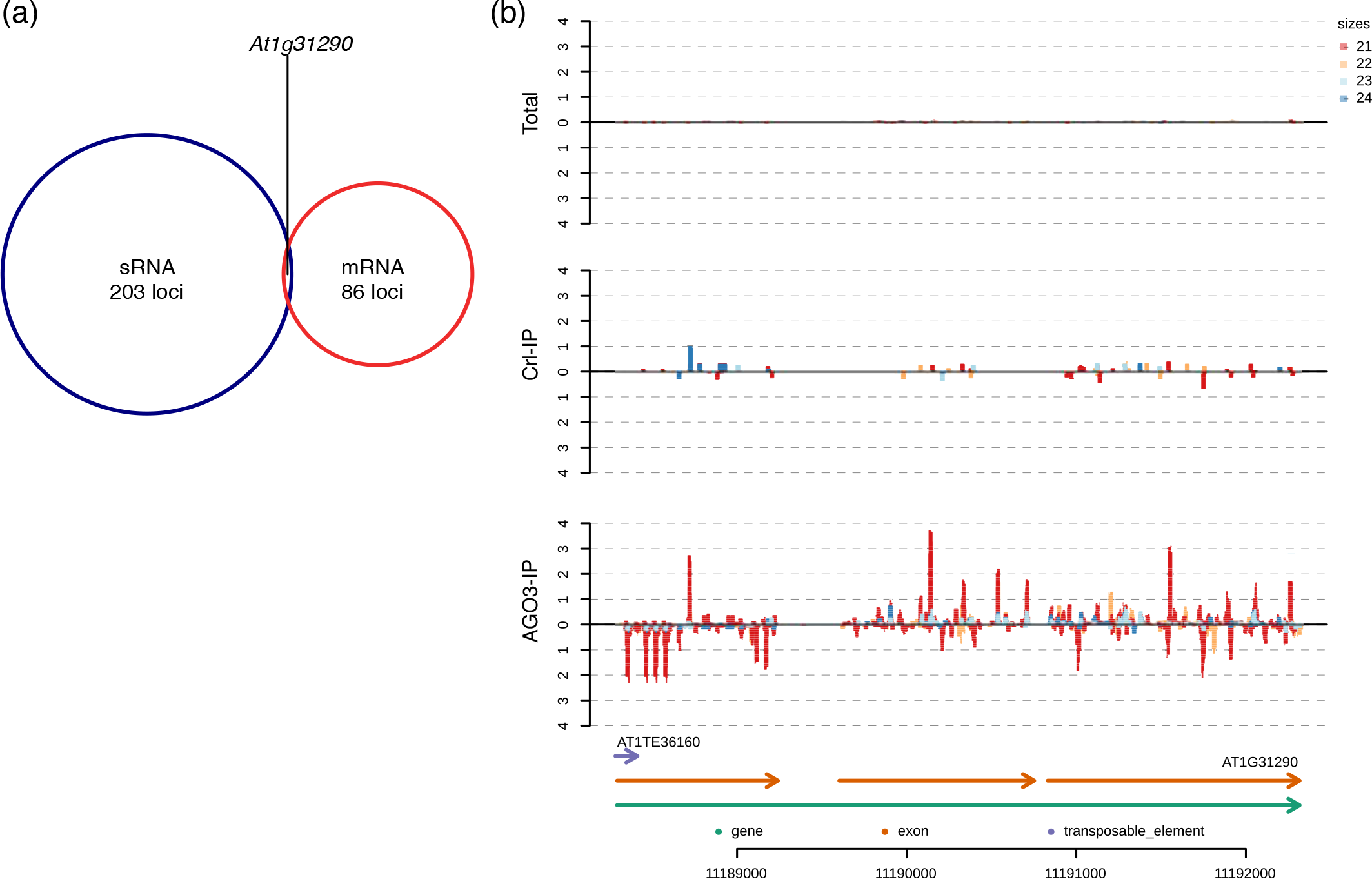
Comparison between AGO3 enriched sRNA and ago3-3 transcriptome. (a) Venn diagram showing a lack of overlap between AGO3 enriched small RNA and loci up or down regulated in ago3-3 mutant. (b) sRNA coverage at the AGO3 locus (At1g31290).

**Figure S9.**
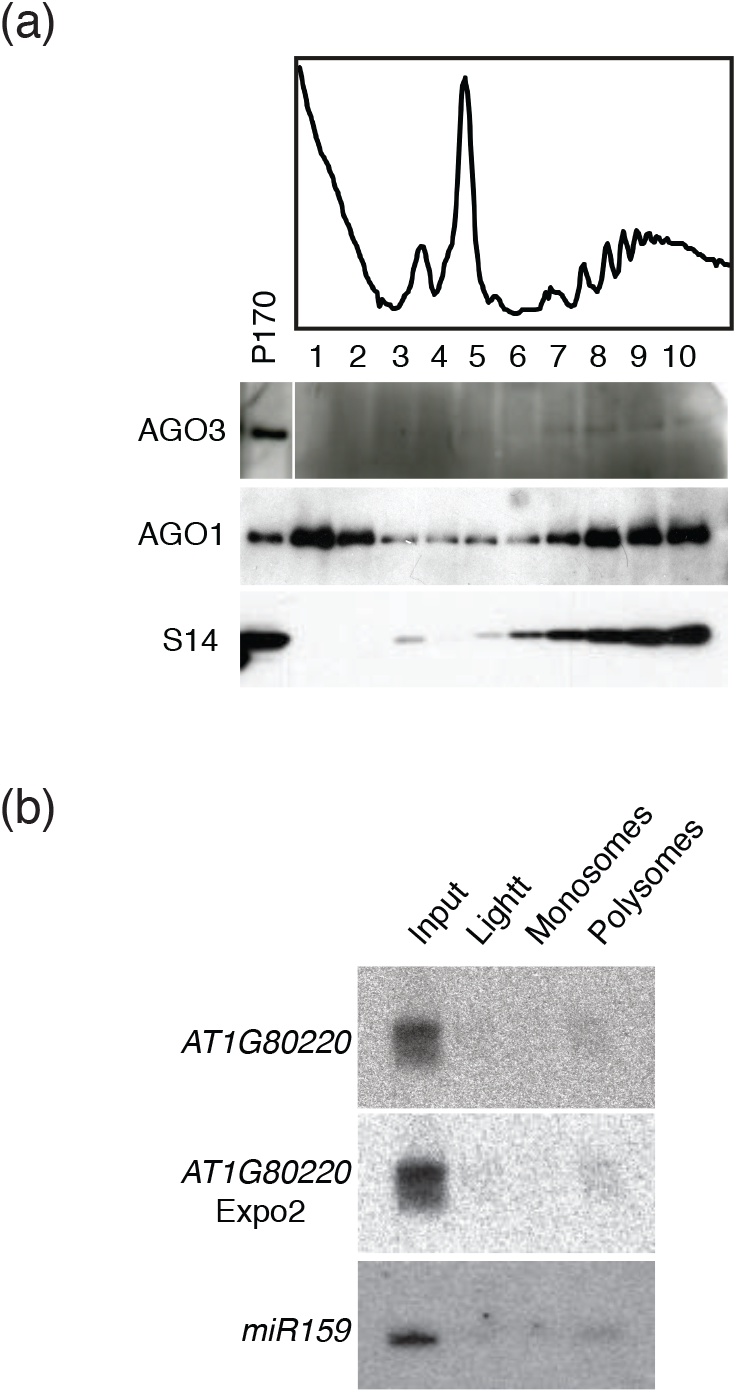
AGO3 and associated small-RNA co-sediments with polysomes. (a) Polysome fractionation showing the co-sedimentation of AGO3 with monosomes and polysomes in 1-5 DAP siliques. P170 represents the fraction loaded onto the sucrose gradient. The ribosomic protein S14 is used as a control to follow the sedimentation of ribosomes within the sucrose gradient. AGO1 is used as a positive control. (b) Northern blot analysis of sRNAs after polysome fractionation on 4-6 DAP siliques. Input represents the raw lysate and miR159 is an internal control for sRNA loading. Light represent a pool of fractions 1-2, Monosomes a pool of fractions 4-5 and Polysomes a pool of fractions 7-9 in reference to Figure 7b.

